# *Cerox1* and microRNA-488-3p noncoding RNAs jointly regulate mitochondrial complex I catalytic activity

**DOI:** 10.1101/323907

**Authors:** Tamara M Sirey, Kenny Roberts, Wilfried Haerty, Oscar Bedoya-Reina, Sebastian Rogatti-Granados, Jennifer Y Tan, Nick Li, Lisa C Heather, Sarah Cooper, Ana C Marques, Chris P Ponting

## Abstract

To generate energy efficiently, the cell is uniquely challenged to co-ordinate the abundance of electron transport chain protein subunits expressed from both nuclear and mitochondrial genomes. How an effective stoichiometry of this many constituent subunits is co-ordinated post-transcriptionally remains poorly understood. Here we show that *Cerox1*, an unusually abundant cytoplasmic long noncoding RNA (lncRNA), modulates the levels of mitochondrial complex I subunit transcripts in a manner that requires binding to microRNA-488-3p. Increased abundance of *Cerox1* cooperatively elevates complex I subunit protein abundance and enzymatic activity, decreases reactive oxygen species production, and protects against the complex I inhibitor rotenone. *Cerox1* function is conserved across placental mammals: human and mouse orthologues effectively modulate complex I enzymatic activity in mouse and human cells, respectively. *Cerox1* is the first lncRNA demonstrated, to our knowledge, to regulate mitochondrial oxidative phosphorylation (OXPHOS) and, with miR-488-3p, represent novel targets for the modulation of complex I activity.

## Introduction

In eukaryotes, coupling of the mitochondrial electron transport chain to oxidative phosphorylation generates the majority of ATP that fulfils cellular energy requirements. The first enzyme of the electron transport chain, NADH:ubiquinone oxidoreductase (complex I), catalyses the transfer of electrons from NADH to coenzyme Q10, pumps protons across the inner mitochondrial membrane and produces reactive oxygen species (ROS). Mammalian mitochondrial complex I dynamically incorporates 45 distinct subunits into a ~1 MDa mature structure[1, 2]. It is known that oxidatively damaged subunits can be exchanged in the intact holo-enzyme[3], but how this process may be regulated is poorly understood. The efficiency and functional integrity of OXPHOS are thought to be partly maintained through a combination of tightly co-ordinated transcriptional and post-transcriptional regulation [4–6] and specific sub-cytoplasmic co-localisation [7, 8]. The nuclear encoded subunits are imported into the mitochondria after translation in the cytoplasm and their complexes assembled together with the mitochondrially encoded subunits in an intricate assembly process [9–11]. Mitochondrial biogenesis is co-ordinated first transcriptionally from both genomes [12], and then post-transcriptionally by regulatory small noncoding RNAs such as microRNAs (miRNAs) [13, 14]. Recently, SAMMSON a long noncoding RNA (lncRNA) was found to bind p32 and, within mitochondria, enhanced the expression of mitochondrial genome-encoded polypeptides [15].

Nuclear-encoded and cytosol-located lncRNAs have not yet been implicated in regulating mitochondrial OXPHOS [16] despite being surprisingly numerous and often found localised to mitochondrion-and ribosome-adjacent portions of the rough endoplasmic reticulum [17]. It is here, on the ribosome, that turnover of miRNA-targeted mRNAs frequently occurs during their translation [18]. Here we describe a novel mammalian conserved lncRNA, termed *Cerox1* (<u>c</u>ytoplasmic <u>e</u>ndogenous <u>r</u>egulator of <u>ox</u>idative phosphorylation 1). *Cerox1* regulates complex I activity in both mouse and human cells by co-ordinately regulating the abundance of at least 12 complex I transcripts via a miRNA-mediated mechanism. *Cerox1* knockdown decreases the enzymatic activities of complexes I and IV. Conversely, elevation of *Cerox1* levels increases their enzymatic activities, halves cellular oxidative stress, and protects cells against the cytotoxic effects of the complex I inhibitor rotenone. To our knowledge, *Cerox1* is the first lncRNA modulator of normal mitochondrial energy metabolism homeostasis and cellular redox state. The miRNA-dependency of *Cerox1* and the regulation of associated OXPHOS transcripts are supported by: (i) direct physical interaction of miR-488-3p with *Cerox1* and complex I transcripts; (ii) decrease or increase in *Cerox1* and complex I transcripts following miR-488-3p overexpression or inhibition, respectively; (iii) miR-488-3p destabilisation of wildtype *Cerox1*, but not a *Cerox1* transcript containing a mutated miR-488-3p MRE seed region; and, (iv) absence of the OXPHOS phenotypes either in cell lines deficient in microRNA biogenesis or when *Cerox1*’s predicted miR-488-3p response element is mutated. The miRNA-dependent role of *Cerox1* illustrates how RNA-interaction networks can regulate OXPHOS and that lncRNAs represent novel targets for modulating OXPHOS enzymatic activity.

## RESULTS

### *Cerox1* is a conserved, ubiquitously expressed long noncoding RNA

*Cerox1* was selected for further investigation from among a set of central nervous system-derived polyadenylated long non-coding RNAs identified by cDNA sequencing (GenBank Accession AK079380, 2810468N07Rik) [19, 20]. Mouse *Cerox1* is a 1.2 kb, two exon, intergenic transcript which shares a bidirectional promoter with the SRY (sex determining region Y)-box 8 (*Sox8*) gene (Fig. 1a). *Cerox1* exons are conserved among eutherian mammals but not with non-eutherian vertebrates (Fig. 1a). A human orthologous transcript (*CEROX1*, GenBank Accession BC098409) was identified by sequence similarity and conserved synteny (60-70% nucleotide identity within alignable regions, Fig. 1b,c). Both mouse and human transcripts have low coding potential (Methods, Supplementary Fig. 1a) and no evidence for translation from available proteomic datasets.

**Figure 1.**
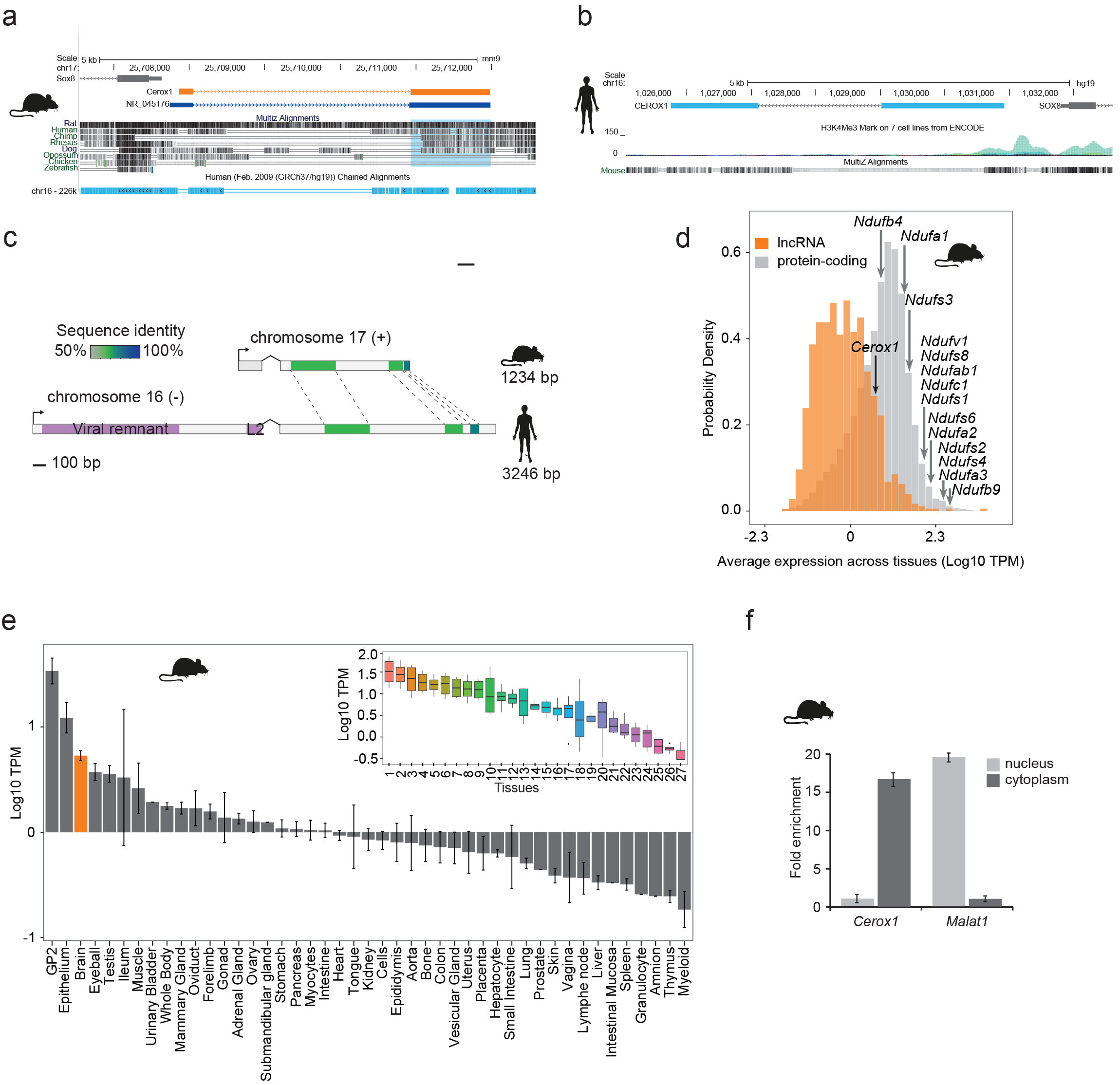
*Cerox1* is an evolutionarily conserved, highly expressed and predominantly cytoplasmic lncRNA. (**a**) The mouse *Cerox1* locus (mm9 assembly). Sequence shaded in blue highlights conservation within exon two among eutherian mammals, but not in non-mammalian vertebrates such as chicken and zebrafish. (**b**) Syntenic human locus (hg19). This transcript was previously identified on the minus strand as *LMF1* non-coding transcript 4, and is located within the intron of *LMF1* non-coding transcript 2. The ENCODE H3K4me3 track (a chromatin mark primarily associated with promoter regions and transcriptional start sites), generated from seven cell lines, reveals peaks in H1-human embryonic stem cells, skeletal muscle, fibroblasts and umbilical vein endothelial cells. (**c**) Schematic representation of mouse *Cerox1* transcript and the human orthologous sequence. Exon two contains blocks of 60-70% sequence identity; human *CEROX1* has an additional 1235 bases of retrotransposed insertions at the 5’ end. (**d**) Distributions of lncRNAs and protein-coding gene expression levels across individuals and tissues in mouse. The black arrow indicates the expression level of *Cerox1*. Expression levels of representative mitochondrial complex I subunits are indicated. TPM = tags per million. (**e**) Average expression levels of *Cerox1* across mouse tissues. The orange bar highlights nervous tissue samples whose values for replicates among neurological tissues are shown in the inset: 1-Medulla oblongata, 2– Spinal cord, 3– Diencephalon, 4– Substantia nigra, 5– Microglia, 6– Raphe, 7– Dorsal spinal cord, 8– Corpora quadrigemina, 9– Cortex, 10– Corpus striatum, 11-Visual cortex, 12– Olfactory brain, 13– Cerebellum, 14– Neurospheres sympathetic neuron derived, 15– Neurospheres parasympathetic neuron derived, 16– Neurospheres enteric neuron derived, 17– Astrocytes (cerebellar), 18– Hippocampus, 19– Hippocampal, 20– Ventral spinal cord, 21– Astrocytes, 22– Pituitary gland, 23– Astrocytes (hippocampus), 24– Cortical neurons, 25– Striatal neurons, 26– Schwann cells, 27– Meningeal cells. Error bars indicate s.e.m. (**f**) Cytoplasmic localisation of mouse *Cerox1* compared to a nuclear retained lncRNA, *Malat1*. Mouse *Cerox1* is 15-fold enriched in the cytoplasm of N2A cells (*n* = 5; error bars s.e.m.).

*Cerox1* expression in primary tissues and cells is exceptionally high, within the top 13% of a set of 879 lncRNAs with associated cap analysis of gene expression (CAGE) clusters (Fig. 1d,e). Expression is particularly high in neuroglia and in neural progenitor cells [21–23], with expression in brain being higher than 64% of all protein coding genes (Fig. 1d). This high expression (Fig. 1e) was confirmed using quantitative real-time PCR (qPCR) for both mouse and human orthologous transcripts (Supplementary Fig. 1b, c). In mouse neuroblastoma (N2A) cells the most highly expressed *Cerox1* transcript (Supplementary Fig. 1d, e) is strongly enriched in the cytoplasmic fraction (Fig. 1f) with a short half-life of 36 ± 16 mins (Supplementary Fig. 1f) and is mainly associated with the ribosome-free fraction (Supplementary Fig. 1g).

### *Cerox1* expression modulates levels of oxidative phosphorylation transcripts

Decreasing or increasing *Cerox1* expression in N2A cells had no effect on the expression of neighbouring genes (Supplementary Fig. 2a). In contrast, *Cerox1* overexpression led to differential expression of 286 distal genes (*q* < 0.05, Bonferroni multiple testing correction; Supplemental Table 1), of which an unexpectedly large majority (83%; 237) were upregulated (*P* < 10^−6^; binomial test). Our attention was immediately drawn to mitochondrial respiratory chain genes because these were considerably (≥20-fold) and significantly enriched among the set of upregulated genes (Fig. 2a).

**Figure 2.**
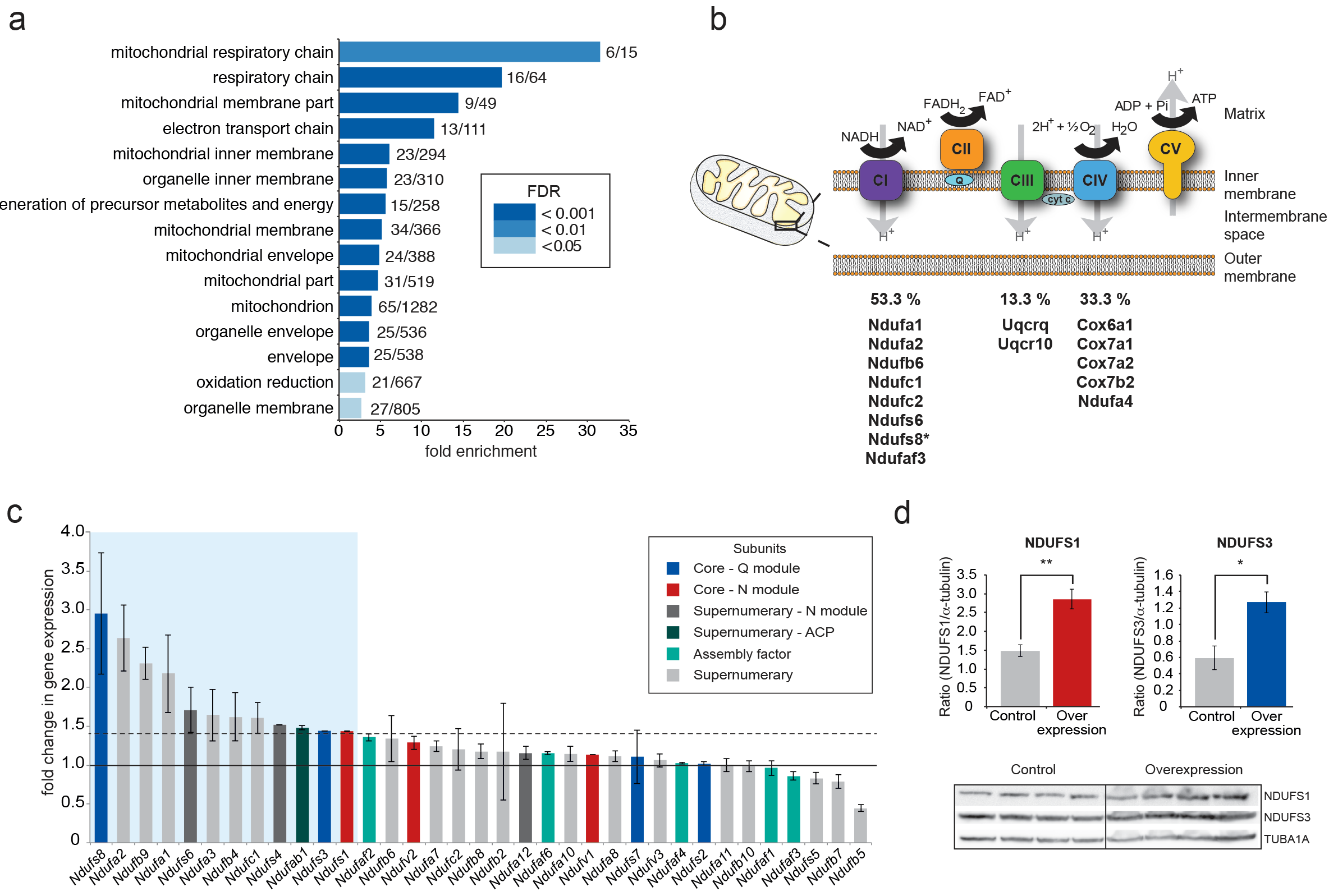
*Cerox1* overexpression elevates levels of OXPHOS transcripts and their encoded proteins. (**a**) Gene ontology analysis indicates a significant enrichment of upregulated genes involved in mitochondrial electron transport, energy production and redox reactions. (**b**) Four membrane bound multi-subunit complexes (CI, CII, CIII, CIV) are embedded in the inner mitochondrial membrane and facilitate transfer of electrons; three of these subunits are also proton pumps which create the chemiosmotic gradient required for ATP synthase activity, with complex V being ATP synthase. The subunits vary in size and complexity with Complex I (NADH:ubiquinone oxidoreductase) consisting of 45 subunits, Complex II (succinate dehydrogenase) 4 subunits, Complex III (Ubiquinol:cytochrome *c* oxidoreductase) 11 subunits and Complex IV (Cytochrome *c* oxidase) 13 subunits. Of 15 oxidative phosphorylation genes whose transcripts were up-regulated following *Cerox1* overexpression 53% were subunits of Complex I, 13% were subunits of Complex III and 33% were subunits of Complex IV. * indicates core subunits that are essential for activity. Note: subunit NDUFA4 has recently been reassigned to mitochondrial complex IV [79]. (**c**) qPCR profiling of 35 complex I subunits and assembly factors (30 nuclear encoded complex I subunits and 5 assembly factors). Transcripts showing a 1.4 fold, or greater, change in expression after overexpression of *Cerox1* are present within the boxed shaded area. The transcripts profiled can be characterised into six categories: Core–Q module, subunits responsible for the electron transfer to ubiquinone; Core–N module, subunits responsible for the oxidation of NADH; Supernumerary subunits– those that are additional to the core subunits required for the catalytic role of complex I, but do not play a catalytic role themselves. Many of these subunits may be performing a structural role, but the majority are of unknown function. The supernumerary subunits can be further subdivided into supernumerary – N module, those accessory subunits associated with the NADH oxidation module of CI; supernumerary ACP (acyl carrier protein) – in addition to being a non-catalytic subunit of CI, NDUFAB1 is also a carrier of the growing fatty acid chain in mitochondrial fatty acid biosynthesis; assembly factor, proteins that are required for the correct assembly and integration of CI. (**d**) Overexpression of *Cerox1* results in large increases in the total protein levels of two core subunits NDUFS1 (average densitometry 1.92, *P* = 0.0043) and NDUFS3 (average densitometry 2.14, *P* = 0.013) for which high quality antibodies exist, normalised to the loading control-tubulin (TUBA1A). NDUFS1 is one of three (NDUFS1, NDUFV1, NDUFV2) core components of the N-module of Complex I. NDUFS3 is one of four (NDUFS2, NDUFS3, NDUFS7, NDUFS8) core components of the Complex I Q-module. Error bars s.e.m. (*n* = 4 biological replicates for control and overexpression). 2-sided *t*-test; ** *P* < 0.01, * *P* < 0.05

The mitochondrial electron transport chain (ETC) consists of five multi-subunit complexes encoded by approximately 100 genes of which only 13 are located in the mitochondrial genome. All 15 ETC transcripts that show statistically significant differential expression after *Cerox1* overexpression are nuclear encoded (Fig. 2b,c) with the greatest changes observed by qPCR for complex I subunit transcripts (Supplementary Fig. 2b). Twelve of 35 nuclear encoded complex I subunits or assembly factors transcripts increased significantly (>40%) following *Cerox1* overexpression; we consider these to be gene expression biomarkers for *Cerox1* activity in the mouse N2A system (Fig. 2c). In the reciprocal *Cerox1* knock-down experiment, all 12 were reduced in abundance, 8 significantly (Supplementary Fig. 2c). Taken together, these results indicate that *Cerox1* positively and co-ordinately regulates the levels of many mitochondrial complex I transcripts.

Increased abundance of OXPHOS subunit transcripts, following *Cerox1* overexpression, was found to elevate protein levels. Western blots using reliable antibodies for the key complex I catalytic core proteins NDUFS1 and NDUFS3, showed approximately 2.0-fold protein level increases that surpassed their ^~^1.4-fold transcript level changes (Fig. 2d). *Cerox1* transcript abundance is thus coupled positively to OXPHOS transcript levels and to their availability for translation, resulting in an amplification of the amount of protein produced.

Cells overexpressing *Cerox1* exhibited a reduction in cell cycle activity, without deviating from normal cell cycle proportions (Supplementary Fig. 2d,e). *Cerox1* levels thus affect the timing of cell division.

### *Cerox1* can regulate mitochondrial OXPHOS enzymatic activity

Increased translation of some complex I transcripts leads to increased respiration [24] or, more specifically, an increase in the enzymatic activity of complex I [25]. Indeed, complex I and complex IV enzymatic activities increased significantly after *Cerox1* overexpression (by 22%, *P* = 0.01; by 30%, *P* = 0.003, respectively; 2-tailed Student’s *t*-test) and oxygen consumption increased (Fig. 3a,b). These moderate increases in enzyme activities are likely to produce substantial increases beyond the already very high basal rate of ATP formation [26]. Conversely, after *Cerox1* knockdown complex I and complex IV enzymatic activities decreased significantly (by 11%, *P* = 0.03 and 19%, *P* = 0.02, respectively), with *Cerox1* knockdown cells consuming less oxygen (Fig. 3c,d). These observed increases in enzymatic activity were not due to changes in mitochondria number because the enzymatic activities of complexes II, III and citrate synthase (Fig. 3a,b), and the mitochondrial-to-nuclear genome ratio all remained unaltered by changes in *Cerox1* levels (Supplementary Fig. 3a, b). These data indicate that *Cerox1* can specifically regulate the catalytic activities of complex I and complex IV in mouse N2A cells.

**Figure 3.**
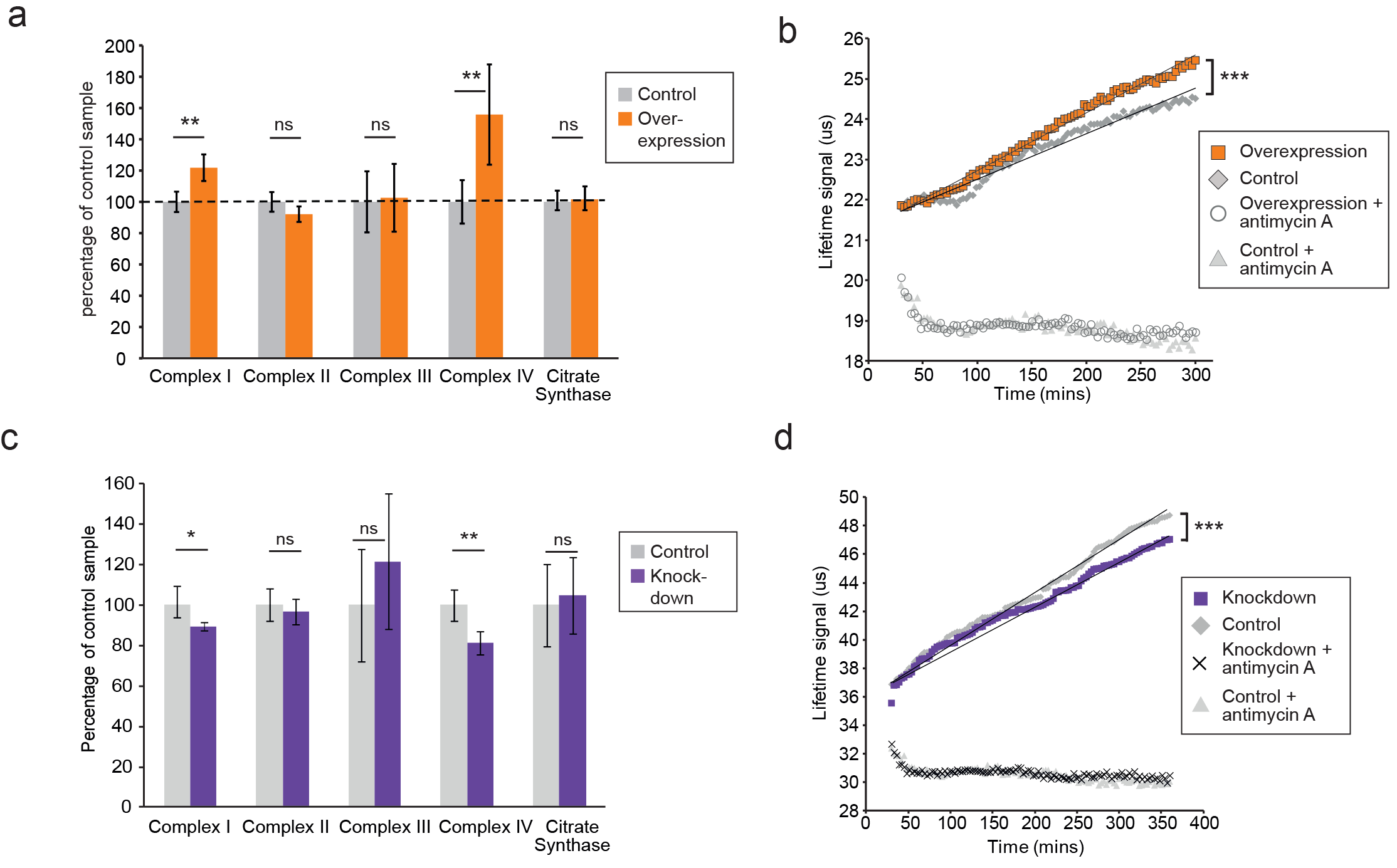
OXPHOS enzyme activity changes concordantly with *Cerox1* level variation. (**a**) Enzyme activities in mouse N2A cells 72 hours post-transfection of *Cerox1* overexpression construct. Mouse *Cerox1* overexpression in N2A cells results in significant increases in the catalytic activities of complexes I (22% increase) and IV (50% increase). Complexes II, III and citrate synthase show no significant change in activity. Error bars s.e.m. (*n* = 8 biological replicates). (**b**) Oxygen consumption, as measured using an extracellular oxygen sensitive probe, by N2A cells overexpressing *Cerox1* over a period of 300 mins. O2 consumption was significantly increased in cells overexpressing *Cerox1* (*P* <0.0001, ANCOVA; Control and Overexpression, *n* = 12 biological replicates, Control + Antimycin A and Overexpression + Antimycin A, *n* = 3 biological replicates). (**c**) shRNA mediated knockdown of *Cerox1* results in significant decreases of Complexes I and IV enzymatic activities 72 hours post transfection; no significant changes were observed for complex III or the citrate synthase control. Error bars s.e.m. (*n* = 8 biological replicates for control and knockdown). (**d**) Oxygen consumption, as measured using an extracellular oxygen sensitive probe, by *Cerox1* knockdown N2A cells over a period of 360 mins. As a negative control, N2A cells were treated with the Complex III inhibitor antimycin A to completely impede the flow of electrons to complex IV. This indicates that the O_2_ consumption measurement specifically reported mitochondrial respiration. O_2_ consumption was significantly decreased in *Cerox1* knockdown cells (*P* = 0.003, ANCOVA; Control and Overexpression, *n* = 12 biological replicates, Control + Antimycin A and Overexpression + Antimycin A, *n* = 3 biological replicates). Enzyme activities are represented as a percentage of control enzyme activity. 2-tailed Student’s *t*-test: ** *P* < 0.01, * *P* < 0.05, ns not significant.

### *Cerox1* expression can protect cells from oxidative stress

Complex I deficient patient cells experience increased ROS production [27]. In *Cerox1* knockdown N2A cells we found ROS levels to have increased significantly, by almost 20% (Fig. 4a). Conversely, in cells overexpressing *Cerox1*, ROS production was nearly halved (Fig. 4a), and protein carbonylation, a measure of ROS-induced damage, was reduced by 35% in *Cerox1*-overexpressing cells (Fig. 4b). The observed *Cerox1*-dependent decrease in ROS levels is of particular interest because mitochondrial complex I is a major producer of ROS which triggers cellular oxidative stress and damage, and an increase in ROS production is a common feature of mitochondrial dysfunction in disease [28].

**Figure 4.**
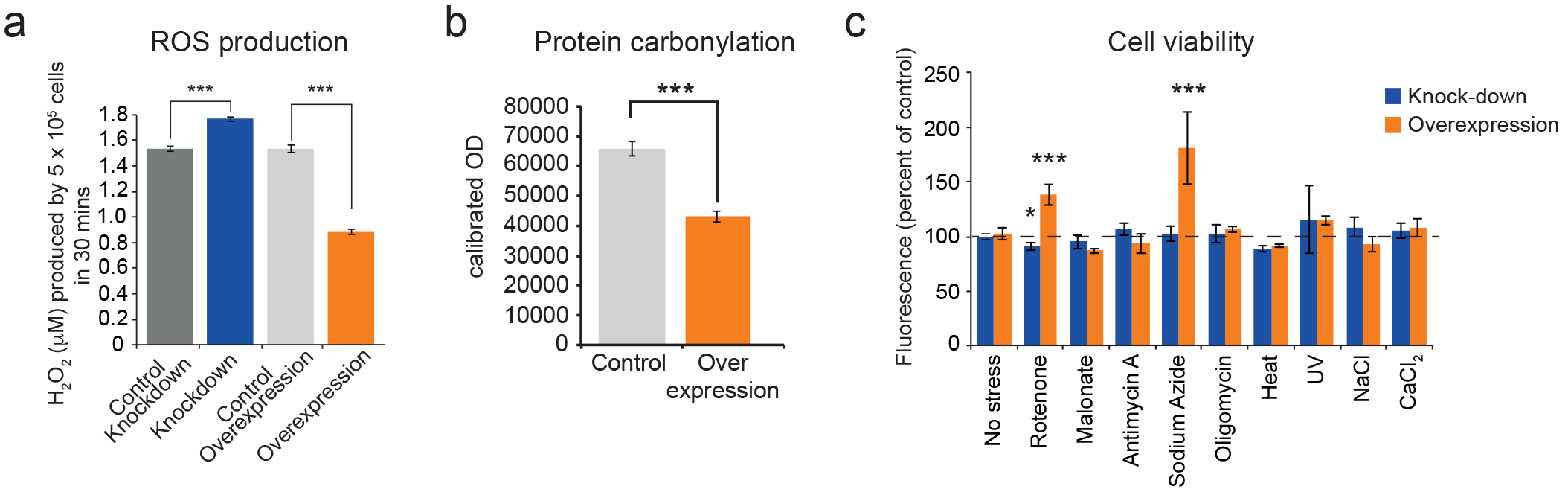
Cellular oxidative stress and viability are *Cerox1* dependent. (**a**) Knockdown of *Cerox1* leads to a 20% increase in the production of reactive oxygen species (*P* = 4.2 × 10^−6^), whilst *Cerox1* overexpression decreases reactive oxygen species production by 45% (*P* = 3.5 × 10^−7^; error bars s.e.m., *n* = 12 biological replicates). (**b**) Protein oxidative damage also decreased in the overexpression condition compared to the control (*P* = 1×10^−3^), as measured by densitometry on western blots against carbonylation of amino acid side chains (error bars s.e.m., *n* = 6 biological replicates). (**c**) Control and *Cerox1* overexpressing and knockdown N2A cells were stressed by the addition of electron transport chain (ETC) inhibitors (rotenone, CI inhibitor; malonate, competitive inhibitor of CII; antimycin A, CIII inhibitor; sodium azide, CIV inhibitor; oligomycin, ATP synthase inhibitor), exposure to environmental stress (heat, ultraviolet radiation), or manipulation of extracellular osmolarity (NaCl) or extracellular calcium (CaCl_2_) concentration, for 1 hour and then the viability of the cells measured using the fluorescent indicator Alamar Blue. Error bars s.e.m. (*n* = 6 biological replicates for control and overexpression). 2-tailed Student’s *t*-test: *** *P* < 0.001, ** *P* < 0.01, * *P* < 0.05.

We next demonstrated that *Cerox1*-induced increases in complex I and complex IV activities protect cells against the deleterious effects of specific mitochondrial complex inhibitors, specifically rotenone and sodium azide (complex I and complex IV inhibitors, respectively); conversely, *Cerox1*-knockdown cells were significantly more sensitive to rotenone (*P* < 1×10^−3^, Fig. 4c). Cells overexpressing *Cerox1* and treated with rotenone, a complex I inhibitor, exhibited no significant difference in protein carbonylation (data not shown). From these results we conclude that increased *Cerox1* expression leads to decreased ROS production, decreased levels of oxidative damage to proteins and can confer protective effects against complex I and complex IV inhibitors.

### Increased OXPHOS enzymatic activity is dependent upon miRNA binding to *Cerox1*

Due to their positive correlation in expression and cytoplasmic localisation we next considered whether *Cerox1* regulates complex I transcripts post-transcriptionally by competing with them for the binding of particular miRNAs. To address this hypothesis, we took advantage of mouse Dicer-deficient (*Dicer*^Δ/Δ^) embryonic stem cells that are deficient in miRNA biogenesis [29]. In wildtype mouse ES cells *Cerox1* overexpression again positively correlated with transcript levels of six complex I subunits (Fig. 5a), of which four had previously shown significant changes in N2A cells after *Cerox1* overexpression (Fig. 2c). In contrast, overexpression in *Dicer*^Δ/Δ^ cells did not lead to an increase in the level of these transcripts (Fig. 5a). These results indicate that *Cerox1*’s ability to alter mitochondrial metabolism is miRNA-dependent.

**Figure 5.**
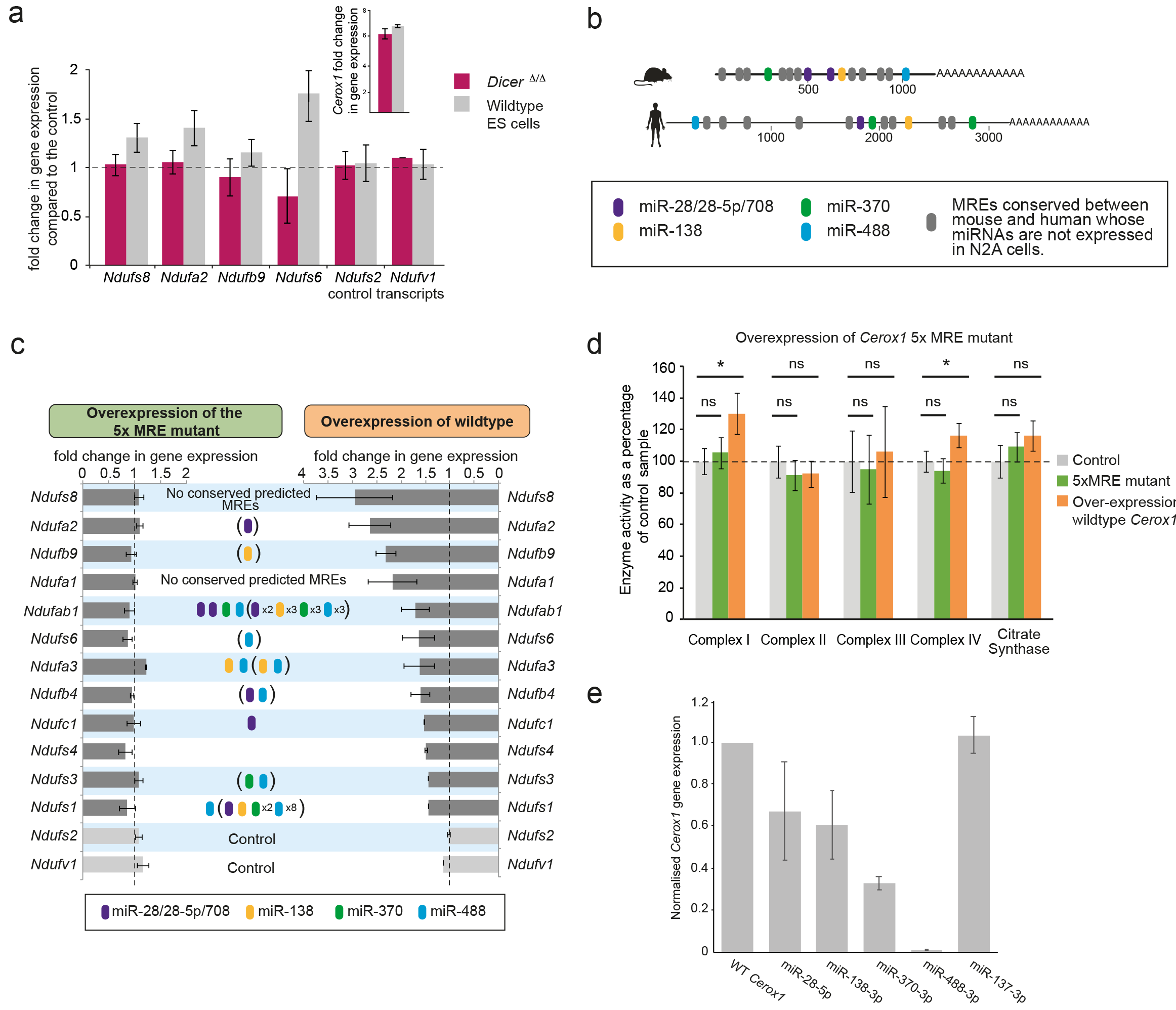
The effect of *Cerox1* on complex I transcript levels is miRNA-dependent. (**a**) Overexpression of *Cerox1* in mouse wildtype and *Dicer* ^Δ/Δ^ embryonic stem (ES) cells (inset graph). The overexpression of *Cerox1* in wildtype mouse embryonic stem cells results in an increase in complex I subunit transcripts, with no observed change in expression of two control subunits (*Ndufs2, Ndufv1)* that were also unaffected in N2A cells. Overexpression of *Cerox1* in *Dicer* ^Δ/Δ^ embryonic stem cells results in no increase in the expression of any complex I subunit. (**b**) Predicted MREs whose presence is conserved in both the mouse and human *Cerox1*. Coloured MREs indicate those MREs whose presence is conserved between mouse and human and whose miRNAs are expressed in N2A cells. miRNA site types are as follows: miR-28-5p, 8mer-A1; miR-138-5p, 6mer; miR-370-3p, 7mer-m8; miR-488-3p-3p, 7mer-m8; miR-708-5p, 7mer-A1. The grey predicted MREs represent those that are conserved, but whose miRNAs are not expressed in N2A cells (miR-125a-3p, miR-199/199-5p, miR-302ac/520f, miR-485/485-5p, miR-486/486-5p, miR-501/501-5p, miR-654-3p, miR-675/675-5p). (**c**) Overexpression of the 5xMRE mutant failed to alter expression of complex I subunit transcripts that otherwise all increase in abundance following wild-type *Cerox1* overexpression. The numbers of MREs predicted by TargetScan v7.0 in these transcripts’ 3’UTRs for the four conserved, N2A expressed miRNA families are indicated (see also Supplemental Table 2). Due to known widespread noncanonical miRNA binding [80], predictions were also extended across the gene body (bracketed MREs). (**d**) Overexpression of the 5xMRE mutant failed to alter OXPHOS enzymatic activity compared to the control for any of the complexes measured. A one-way ANOVA was applied to test for differences in activities of the mitochondrial complexes between a control and overexpression of wildtype *Cerox1* and the 5xMRE mutant. A post-hoc Dunnett’s test indicated that the overexpression of wildtype *Cerox1* resulted, as expected, in significantly increased complex I and IV activities (*F* [2, 21] = 4.9, *P* = 0.017; *F*[2, 20] = 4.6, P = 0.033), while comparisons for the 5xMRE mutant with the control were not significant. There was no significant difference in the activities of complex III (*F*[2,19]= 0.08, *P* = 0.5) or citrate synthase (*F*[2,20]=2.6, *P* = 0.42). Significance levels, one-way ANOVA, Dunnett’s post hoc test * *P* < 0.05. **(e**) Four to six fold overexpression of each of four miRNAs with predicted MREs whose presence is conserved in both mouse and human *Cerox1* resulted in a decrease in *Cerox1* transcript level, with overexpression of miR-488-3p resulting in >90% knock down of *Cerox1*. This was not observed when the miRNA miR-137-3p, which has no predicted MREs in *Cerox1*, was overexpressed.

Four specific miRNA families (miR-138-5p, miR-28/28-5p/708-5p, miR-370-3p, and miR-488-3p; Supplementary Fig. 4a) were selected for further investigation based on the conservation of their predicted binding sites (miRNA recognition elements, MREs) in both mouse *Cerox1* and human *CEROX1* (Fig. 5b). All MREs conserved in mouse *Cerox1* and human *CEROX1* for N2A-expressed miRNAs (Fig. 5b) were mutated by inversion of all five seed regions. This mutated *Cerox1* transcript failed to alter either complex I transcript levels or enzyme activities when overexpressed (Fig. 5c, d) indicating that its molecular effects are mediated by one or more of these MREs.

We then overexpressed each of the four miRNAs in N2A cells which each caused a reduction in *Cerox1* levels (Fig. 5e). Overexpression of the tissue-restricted miRNA miR-488-3p (Supplementary Fig. 4b, [30]) resulted in the greatest depletion of the *Cerox1* transcript (Fig. 5e), indicating that this MRE is likely to be physiologically relevant[31].

### *Cerox1* activity is mediated by miR-488-3p

Our previous results showed that *Cerox1* abundance modulates complex I activity and transcripts (Figs. 2–4) and that miR-488-3p has the greatest effect in decreasing *Cerox1* transcript levels (Fig. 5e). To determine whether miR-488-3p modulates complex I transcript levels we overexpressed and inhibited miR-488-3p in N2A cells (Fig. 6a,b). Results showed that miR-488-3p modulates these transcripts’ levels, with overexpression leading to a significant downregulation of all 12 *Cerox1*-sensitive complex I transcripts (Fig. 6a), whilst, conversely, miR-488-3p inhibition leads to increased expression (Fig. 6b).

**Figure 6.**
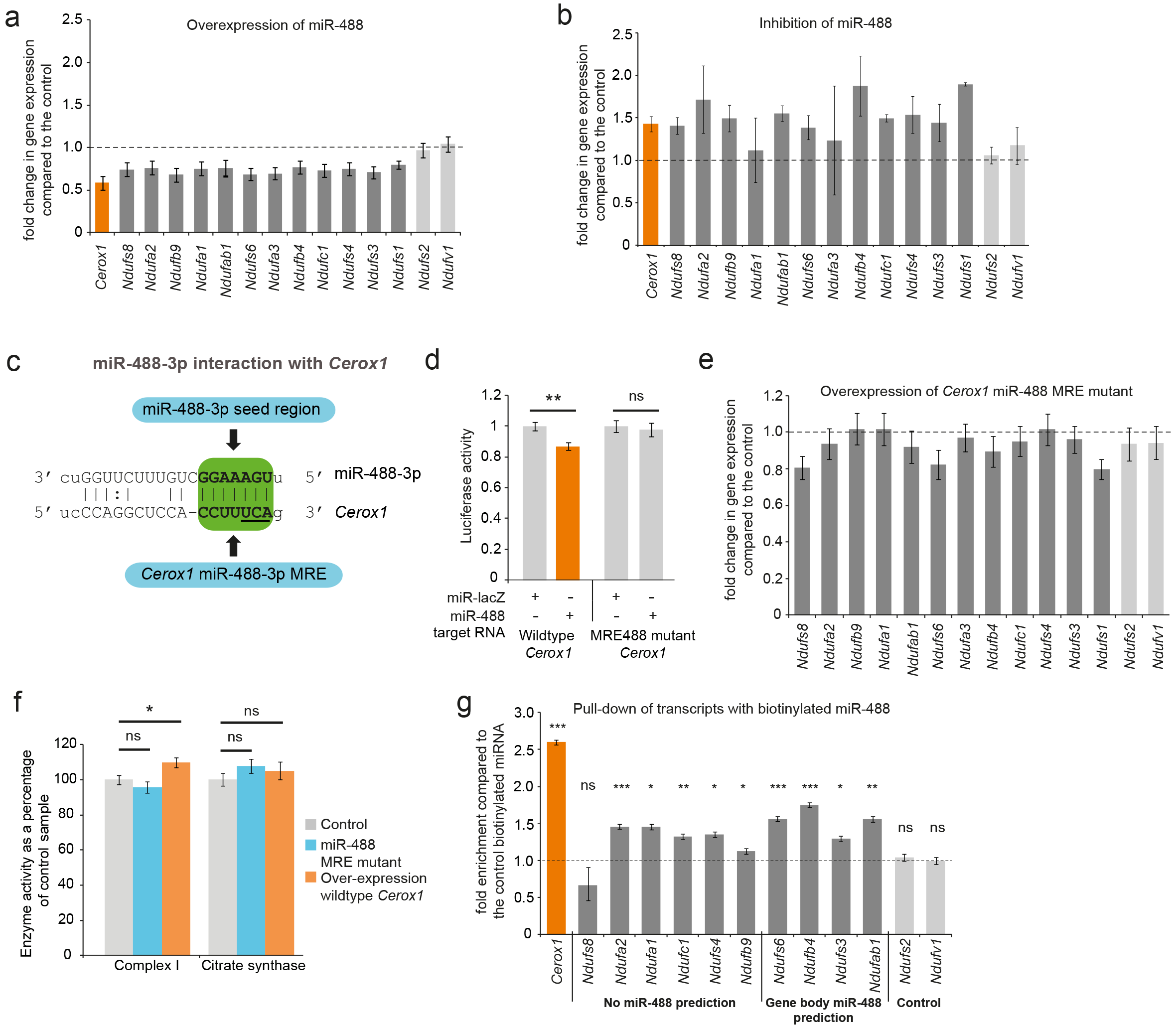
An intact miR-488-3p response element site is required for the effect of *Cerox1* on complex I catalytic activity. (**a**) Overexpression of miR-488-3p knocks down all *Cerox1*-sensitive subunit transcripts. (**b**) Inhibition of miR-488-3p increases the expression of most (11/13) target transcripts compared to the control. (**c**) Schematic of the predicted miR-488-3p miRNA recognition element in *Cerox1.* The interaction of miR-488-3p with *Cerox1* is predicted to involve a 7mer-8m seed site, with the heptamer sequence of the seed being complementary to nucleotides 2-8 of the miRNA. Underlined residues indicate the location of the seed region mutation. (**d**) Luciferase destabilisation assay for both wildtype *Cerox1* and *Cerox1* mutated within the miR-488-3p MRE. (**e, f**) Overexpression of *Cerox1* mutated within a single miR-488-3p MRE (**e**) failed to alter expression levels of complex I subunits that increase in expression with wild-type *Cerox1* overexpression and (**f**) failed to recapitulate the increase in complex I catalytic activity observed for the wildtype transcript. As expected, wildtype enzymatic activity was significantly different for complex I (*F* [2, 19] = 4=8.8, *P* = 0.019). A post-hoc Dunnett’s test indicated that the overexpression of wildtype *Cerox1* resulted in significantly increased complex I activity, while the comparison of the *Cerox1* miR-488-3p MRE mutant with the control was not significant. There was no significant difference in the activity of citrate synthase (*F*[2,21]=1.4, *P* = 0.28). Significance levels, one-way ANOVA, Dunnett’s post hoc test * *P* < 0.05. **(g)** Enrichment of 9 *Cerox*1 sensitive transcripts that do not have predicted canonical miR-488-3p MREs using biotinylated miR-488-3p as bait as compared to the control biotinylated miRNA. 2-sided *t*-test; *** *P* < 0.001, ** *P* < 0.01, * *P* < 0.05, ns – not significant.

To determine whether the single predicted miR-488-3p MRE in *Cerox1* is required to exert its effects on complex I we created a *Cerox1* transcript containing three mutated nucleotides in this MRE (Fig. 6c). As expected for a bona fide MRE, these substitutions abrogated the ability of miR-488-3p to destabilise *Cerox1* transcript (Fig. 6d). Importantly, these substitutions also abolished the ability of *Cerox1*, when overexpressed, to elevate complex I transcript levels (Fig. 6e), and to enhance complex I enzymatic activity (Fig. 6f). The latter observation is important because not all bona fide miRNA-transcript interactions are physiologically active [31]. Finally, direct interaction between *Cerox1* and miR-488-3p was confirmed by pulling-down transcripts with biotinylated miR-488-3p (Fig. 6g). This experiment also identified 9 of 10 complex I transcripts tested as direct targets of miR-488-3p binding. These included transcripts not predicted as containing a miR-488-3p MRE, as expected from the high false negative rate of MRE prediction algorithms [32–34]. As expected, the two negative control transcripts, which are not responsive to *Cerox1* transcript levels, and have no predicted MREs for miR-488-3p failed to bind miR-488-3p.

Considered together, these findings indicate that: (i) *Cerox1* can post-transcriptionally regulate OXPHOS enzymatic activity as a miRNA decoy, and (ii) of 12 miR-488-3p:*Nduf* subunit interactions that were investigated, all 12 are indicated either by responsiveness to miR-488-3p through miRNA overexpression or inhibition (Fig. 6a,b), or by direct interaction with a biotinylated miR-488-3p mimic (Fig. 6f). Consequently, our data demonstrates that miR-488-3p directly regulates the transcript levels of *Cerox1* and at least 12 nuclear encoded mitochondrial complex I subunit genes (31% of all) and indirectly modulates complex I activity (Fig. 6h) in N2A cells.

### *Cerox1* is an evolutionarily conserved regulator of mitochondrial complex I activity

Fewer than 20% of lncRNAs are conserved across mammalian evolution [35] and even for these functional conservation has rarely been investigated. In our final set of experiments we demonstrated that *CEROX1*, the orthologous human transcript, is functionally equivalent to mouse *Cerox1* in regulating mitochondrial complex I activity. Similar to mouse *Cerox1,* human *CEROX1* is highly expressed in brain tissue, is otherwise ubiquitously expressed (Supplementary Fig. 1b), and is enriched in the cytoplasm (Fig. 7a). *CEROX1* is expressed in human tissues at unusually high levels: it occurs among the top 0.3% of all expressed lncRNAs (Fig. 7b) and its average expression is higher than 87.5% of all protein coding genes [36]. Its expression is highest within brain tissues, particularly within the basal ganglia and cortex (Fig. 7c).

**Figure 7.**
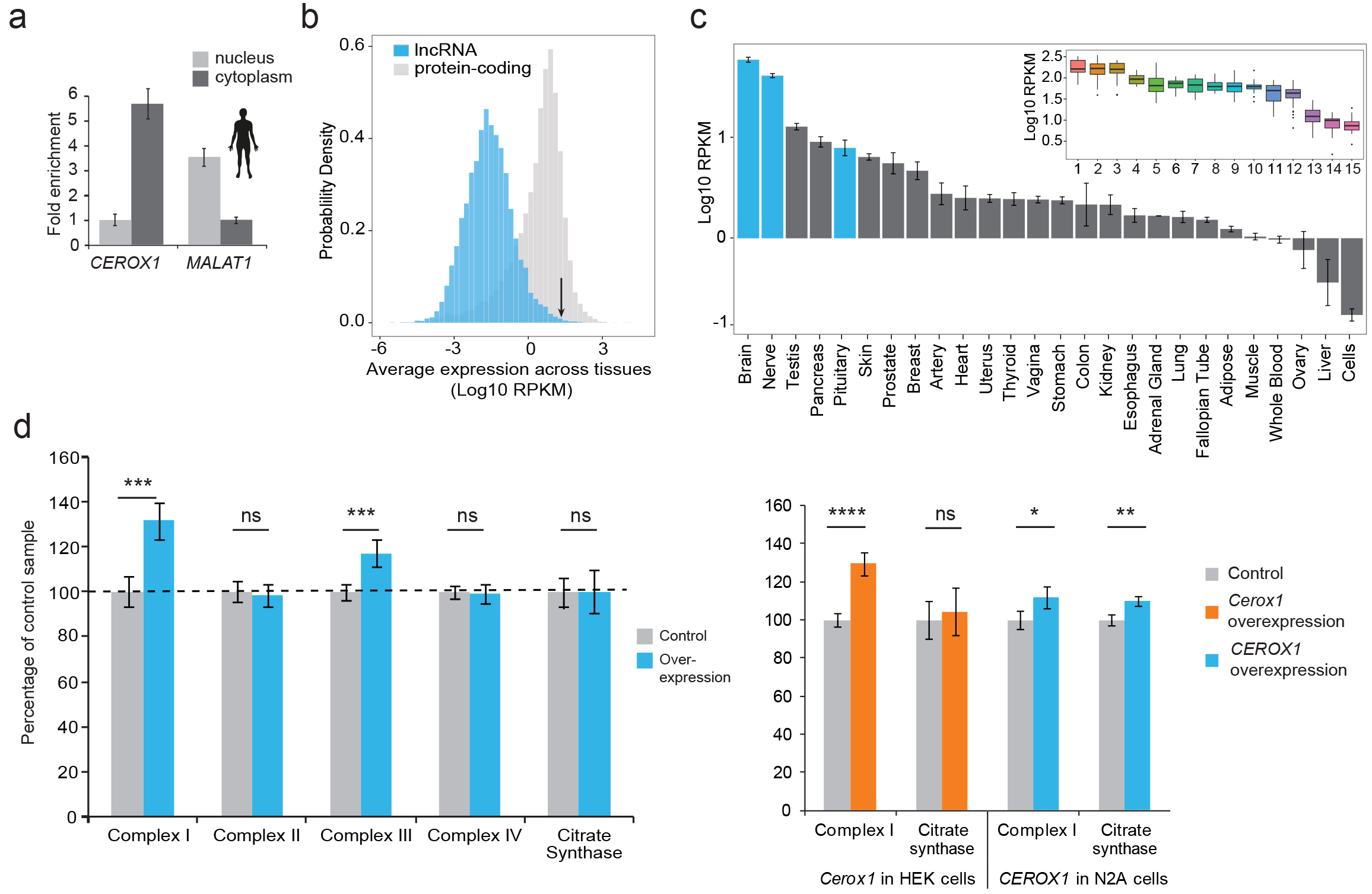
Human *CEROX1* modulates complex I activity in mouse cells. (**a**) *CEROX1* is enriched in the cytoplasm. (**b**) Relative levels of lncRNA (blue) and protein-coding gene (grey) expression across individuals and tissues in human. The black arrow indicates the expression level of *CEROX1* in the set of 5161 lncRNAs. RPKM: reads per kilobase per million reads. (**c**) Average expression levels of *CEROX1* in human tissues. Blue bars highlight neurological tissues used to build the inset graph. The inset graphic represents the comparison of gene expression variation among individuals for neurological tissues: 1–Putamen, 2-Caudate nucleus, 3-Nucleus accumbens, 4–Cortex, 5-Substantia nigra, 6–Amygdala, 7–Hippocampus, 8-Spinal cord, 9-Anterior cingulate cortex, 10-Frontal cortex, 11–Hypothalamus, 12-Tibial nerve, 13–Cerebellum, 14–Pituitary gland, 15-Cerebellar hemisphere. (**d**) OXPHOS enzyme activities in human HEK293 cells after 72 hours of *CEROX1* overexpression. Overexpression of *CEROX1* results in significant increases in the activities of complexes I (31% increase) and III (18% increase), with no significant change in other enzyme activities. Error bars s.e.m. (*n* = 8 biological replicates). *t*-test: *** *P* < 0.001, ns=not significant. (**e**) Reciprocal overexpression of mouse *Cerox1* in human HEK293 cells or human *CEROX1* in mouse N2A cells results in elevated complex I activity. Error bars s.e.m. (*n* = 8 biological replicates). *t*-test: **** *P* <0.0001, ** *P* < 0.01, * *P* <0.05, ns=not significant.

Importantly, following *CEROX1* overexpression in human embryonic kidney (HEK293T) cells mitochondrial complexes’ I and III activities increased significantly (Fig. 7d). *CEROX1* overexpression had a greater effect on complex I activity than the mouse orthologous sequence and also increased the activity of complex III, rather than complex IV activity, in these cells. The latter distinction could reflect the differences in miRNA pools between mouse and human cell lines and/or the presence of different MREs in the lncRNA and human OXPHOS transcripts.

Strikingly, either reciprocal expression of mouse *Cerox1* in human HEK293T cells or human *CEROX1* in mouse N2A cells, recapitulates the previously observed increase in complex I activity (Fig. 7e). The role of both *Cerox1* and *CEROX1* in modulating the activity of mitochondrial complex I has thus been conserved over 90 million years since the last common ancestor of mouse and human.

## DISCUSSION

*Cerox1* is the first evolutionarily conserved lncRNA to our knowledge that has been demonstrated experimentally to regulate mitochondrial energy metabolism. Its principal location in N2A cells is in the cytoplasm where it post-transcriptionally regulates the levels of mitochondrial OXPHOS subunit transcripts and proteins by decoying for miRNAs (Supplementary Fig. 5), most particularly miR-488-3p. This microRNA shares with *Cerox1* an early eutherian origin, and enhanced expression in brain samples [30]. Changes in *Cerox1* abundance *in vitro* alter mitochondrial OXPHOS subunit transcript levels and, more importantly, elicit larger changes in their protein subunits levels, leading to substantial changes in mitochondrial complex I catalytic activity. The observed changes in catalytic activity are in line with the degree of change seen in diseases exhibiting mitochondrial dysfunction for example [37–39]. Overexpression of *Cerox1* in N2A cells increases oxidative metabolism, halves cellular oxidative stress and enhances protection against the complex I inhibitor rotenone. The effect of *Cerox1* on complex I subunit transcript levels can be explained by their sharing MREs with *Cerox1*, and subsequent competition for miRNA binding, most notably for miR-488-3p, which buffers the OXPHOS transcripts against miRNA-mediated repression.

Multiple RNA transcripts have been experimentally shown to compete with mRNAs for binding to miRNAs, thereby freeing the protein coding mRNA from miRNA-mediated repression [40–45]. It has been experimentally demonstrated that this miRNA:RNA regulatory crosstalk can initiate rapid co-ordinate modulation of transcripts whose proteins participate within the same complex or process [45]. Physiological relevance of this crosstalk mechanism remains incompletely understood. Furthermore, mathematical modelling [46–48] and experimental investigation [49, 50] of the dynamics and mechanism of endogenous transcript competition for miRNA binding have resulted in contrasting conclusions. Current mathematical models do not take full account of miRNA properties, such as the repressive effect not being predictable from its cellular abundance [51], intracellular localisation such as at the rough ER [52], loading on the RNA-induced silencing complex (RISC) (Flores et al., 2014), or AGO2’s phosphorylation status within the RISC (Golden et al. 2017). The conclusions of experiments have also assumed that all miRNA target transcripts that contain the same number and affinity of miRNA binding sites are equivalent, that steady-state measurements are relevant to repression dynamics, and that observations for one miRNA in one experimental system are equally applicable to all others [53].

Considered together, our lines of experimental evidence indicate that miRNA-mediated target competition by *Cerox1* substantially perturbs a post-transcriptional gene regulatory network that includes at least 12 complex I subunit transcripts. This is consistent with the expression level of miR-488-3p [54] and the high *in vivo* expression of both *Cerox1* and OXPHOS transcripts [55, 56]. Human *CEROX1* levels, for example, exceed those of all complex I subunit transcripts (those in Fig. 6a) in both newly-formed and myelinating oligodendrocytes [57]. High levels of *Cerox1* in the central nervous system support the notion that *Cerox1* maintains OXPHOS homeostasis in cells with sustained high metabolic activity and high energy requirements [58].

Our experiments demonstrate that post-transcriptional regulation of a subset of complex I subunits by *Cerox1* leads to elevated oxygen consumption. How consumption increases when there is a higher abundance of only a subset of OXPHOS transcripts remains unclear. However, this phenomenon has been observed previously in mouse dopaminergic neurons [25] and primary mouse embryonic fibroblasts and pinnal tissues [24]. Our observation of increased enzymatic activity may relate to the formation, by the complexes of the respiratory chain, of higher order supercomplexes [59, 60]. Alternatively, the observed increases in OXPHOS activity may reflect some subunits of the complex I holo-enzyme (including NDUFS3 and NDUFA2) being present as a monomer pool and therefore being available for direct exchange without integration into assembly intermediates [3]. This monomer pool facilitates the rapid swapping out of oxidatively damaged complex I subunits [3]. It is thus possible that *Cerox1*-mediated expansion of the monomer pool thereby improves complex I catalysis efficiency (Supplemental Fig. 5).

More efficient ETC enzymatic activity might be relevant to mitochondrial dysfunction, a feature of many disorders that often manifests as decreases in the catalytic activities of particular mitochondrial complexes. A decrease in catalytic activity can result in elevated ROS production, leading to oxidative damage of lipids, DNA, and proteins, with OXPHOS complexes themselves being particularly susceptible to such damage [61]. Parkinson’s and Alzheimer’s diseases both feature pathophysiology associated with oxidative damage resulting from increased ROS production and decreased complex I and IV activities (a reduction of 30% and 40%, respectively) [37, 62, 63]. Currently no effective treatments exist that help to restore mitochondrial function. It has been demonstrated that a 20% increase in complex I activity protects mouse midbrain dopaminergic neurons against MPP+, a complex I inhibitor and a chemical model of Parkinson’s disease [25]. We note that highest expression of *CEROX1* occurs in the basal ganglia, which contains the substantia nigra in which the dopaminergic neurons that are particularly sensitive to degeneration in Parkinson’s disease are located. *CEROX1*’s ability to increase mitochondrial complex I activity might be recapitulated pharmacologically to restore mitochondrial function, as an exemplar of therapeutic upregulation of gene expression [64].

## MATERIALS AND METHODS

### Gene expression profiling

The lncRNA transcripts were assessed for coding potential using the coding potential calculator [65], PhyloCSF [66] and by mining proteomics and small open reading frame resources for evidence of translation [67–69].

5’ and 3’ ends of the mouse and human lncRNA transcripts were confirmed by 5’ and 3’ RACE using the GeneRacer™ Kit (Invitrogen) according to the manufacturer’s instructions. Total RNA from twenty normal human tissues (adipose, bladder, brain, cervix, colon, oesophagus, heart, kidney, liver, lung, ovary, placenta, prostate, skeletal muscle, small intestine, spleen, testes, thymus, thyroid and trachea) were obtained from FirstChoice® Human Total RNA Survey Panel (Invitrogen). Total RNA from twelve mouse tissues (bladder, brain, colon, heart, kidney, liver, pancreas, skeletal muscle, small intestine, stomach and testis) were obtained from Mouse Tissue Total RNA Panel (Amsbio). RNA from cell lines was extracted using the RNeasy mini kit (Qiagen) according to the manufacturer’s instructions, using the optional on column DNase digest. cDNA synthesis for all samples was performed on 1 μg of total RNA using a QuantiTect Reverse Transcription kit (Qiagen) according to the manufacturer’s instructions. RNA was extracted from samples used for the detection of miRNAs using the miRNeasy mini kit (Qiagen) according to the manufacturer’s instructions (with on column DNase digest). All RNA samples were quantified using the 260/280 nm absorbance ratio, and RNA quality assessed using a Tapestation (Agilent). RNA samples with an RNA integrity number (RIN) >8.5 were reverse transcribed. 1 μg of total RNA from the miRNA samples were reverse transcribed using the NCode VILO miRNA cDNA synthesis kit. Expression levels were determined by real-time quantitative PCR, using SYBR® Green Master Mix (Applied Biosystems) and standard cycling parameters (95° 10 min; 40 cycles 95° 15s, 60° 1 min) followed by a melt curve using a StepOne™ thermal cycler (Applied Biosystems). All amplification reactions were performed in triplicate using gene specific primers. Multiple reference genes were assessed for lack of variability using geNorm [70]. Human expression data were normalised to *TUBA1A* and *POLR2A*, whilst mouse expression data were normalised to *Tbp* and *Polr2a*.

### Tissue culture and flow cytometry

Mouse Neuro-2a neuroblastoma cells (N2A) and human embryonic kidney (HEK293T) cells were bgrown at 37° in a humidified incubator supplemented with 5% CO_2_. Both cell lines were grown in Dulbecco’s modified Eagle medium containing penicillin/streptomycin (100 U/ml, 100 ug/ml respectively) and 10% fetal calf serum. Cells were seeded at the following densities: 6 well dish, 0.3 × 10^6^; 48 well dish, 0.2 × 10^4^; T75 flask 2.1 × 10^6^. Mouse embryonic stem cells and dicer knock-out embryonic stem cells were maintained as described previously [29]. Cells were counted using standard haemocytometry. For flow cytometry the cells were harvested by trypsinization, washed twice with PBS and fixed in 70% ethanol (filtered, −20°). The cell suspension was incubated at 4° for 10 min and the cells pelleted, treated with 40 μg/ml RNase A and propidium iodide (40 μg/ml) for 30 min at room temperature. Cells were analysed using a FACSCalibur (BD-Biosciences) flow cytometer.

### Transcript localisation and RNA turnover

Cells were fractionated into nuclear and cytoplasmic fractions in order to determine the predominant cellular localization of lncRNA transcripts. Briefly, approximately 2.8 × 10^6^ cells were collected by trypsinization, washed three times in PBS and pelleted at 1000 g for 5 min at 4°C. The cell pellet was resuspended in 5 volumes of lysis buffer (10 mM Tris-HCl, pH 7.5, 3 mM MgCl_2_, 10 mM NaCl, 5 mM EGTA, 0.05% NP40, and protease inhibitors [Roche, complete mini]) and incubated on ice for 15 min. Lysed cells were then centrifuged at 2000 g for 10 min at 4°C, and the supernatant collected as the cytoplasmic fraction. Nuclei were washed three times in nuclei wash buffer (10 mM HEPES, pH 6.8, 300 mM sucrose, 3 mM MgCl_2_, 25 mM NaCl, 1 mM EGTA), and pelleted by centrifugation at 400 g, 1 min at 4°C. Nuclei were extracted by resuspension of the nuclei pellet in 200 μl of nuclei wash buffer containing 0.5% Triton X-100 and 700 units/ml of DNase I and incubated on ice for 30 mins. Nucleoplasm fractions were collected by centrifugation at 17 000g for 20 min at 4°. RNA was extracted as described above, and RNA samples with RIN values >9.0 used to determine transcript localisation.

To determine the stability of the lncRNA transcripts, cells were cultured to ^~^50 % confluency and then transcription was inhibited by the addition of 10 μg/ml actinomycin D (Sigma) in DMSO. Control cells were treated with equivalent volumes of DMSO. Transcriptional inhibition of the N2A cells was conducted for 16 hours with samples harvested at 0 hrs, 30 mins, 1 hr, 2 hrs, 4 hrs, 8 hrs and 16 hrs. RNA samples for fractionation and turnover experiments were collected in Trizol (Invitrogen) and RNA purified and DNAse treated using the RNeasy mini kit (Qiagen). Reverse transcription for cellular localisation and turnover experiments was performed as described earlier.

### Constructs and biotinylated miRNAs

PCR primers modified to contain BglII and XhoI sites were used to amplify the full length mouse *Cerox1*, whilst human *CEROX1* and the mouse 5x MRE mutant were synthesised by Biomatik (Cambridge, Ontario), and also contained BglII and XhoI sites at the 5’ and 3’ ends respectively. All other MRE mutants were produced using overlapping PCR site directed mutagenesis to mutate 3 bases of the miRNA seed region. All purified products were ligated into the prepared backbone and then transformed by heat shock into chemically competent DH5α, and plated on selective media. All constructs were confirmed by sequencing. All full length lncRNAs were cloned into the pCAGGS overexpression vector under the βactin/β-globin promoter. As an overexpression/transfection control EGFP was cloned into the pCAGGS backbone. Short hairpin RNAs specific to the transcripts were designed using a combination of the RNAi design tool (Invitrogen) and the siRNA selection program from the Whitehead Institute [71]. Six pairs of shRNA oligos to the target genes and β-galactosidase control oligos were annealed to create double-stranded oligos and cloned into the BLOCK-iT™ U6 vector (Invitrogen), according to the manufacturer’s instructions. miRNA expression constructs were generated and cloned into the BLOCK-iT™ Pol II miR RNAi expression vector (Invitrogen) according to the manufacturer’s instructions. miRNA inhibitors were sourced from Ambion and used according to manufacturer’s instructions.

One day prior to transfection cells were either seeded in 6 well dishes (0.3 × 10^6^ cells/well), or in T75 flasks (2.1 × 10^6^ cells/flask). Twenty-four hours later cells in 6 well dishes were transfected with 1 μg of shRNA, miRNA or overexpression construct and their respective control constructs using FuGENE® 6 (Promega) according to the manufacturer’s guidelines. Cells in T75 flasks were transfected with 8 μg of experimental or control constructs. Transfected cells were grown for 48 hours under standard conditions, and then harvested for either gene expression studies or biochemical characterisation. Efficacy of the overexpression and silencing constructs was determined by real-time quantitative PCR.

Transcripts for the luciferase destabilisation assays were cloned into the pmirGLO miRNA target expression vector (Promega) and assayed using the dual-luciferase**®** reporter assay system (Promega). miRCURY LNA biotinylated miRNAs (mmu-miR-488-3p and mmu-negative control 4) were purchased from Exiqon, and direct mRNA-miRNA interactions were detected using a modified version of [72] and enrichment of targets was detected by qPCR.

### Computational techniques

MREs were predicted using TargetScan v7.0 in both the 3’UTR (longest annotated UTR, ENSEMBL build 70) or the full length transcript of protein coding genes, and across the entire transcript for lncRNAs. The average expression across 46 human tissues and individuals according to the Pilot 1 data from the GTEx Consortium [73] was computed for both protein-coding genes and intergenic lncRNAs from the Ensembl release 75 annotation [74]. We used the normalized number of CAGE tags across 399 mouse cells and tissues from the FANTOM5 Consortium (http://fantom.gsc.riken.jp)[75] as an approximation of expression levels for protein-coding genes and intergenic lncRNAs from the Ensembl release 75 annotation. If multiple promoters were associated with a gene, we selected the promoter with the highest average tag number. Conserved sequence blocks in the lncRNA sequences were identified using LALIGN [76].

### Microarray analysis

Microarray analysis was performed on 16 samples (four overexpression/four overexpression controls; four knock-down/four knock-down controls), and hybridizations were performed by the OXION array facility (University of Oxford). Data were analysed using the web-based Bioconductor interface, CARMAweb [77]. Differentially expressed genes (Bonferroni corrected *P*-value <0.05) were identified between mouse lncRNA overexpression and control cells using Limma from the Bioconductor package between the experimental samples and the respective controls. Microarray data are accessible through ArrayExpress, accession E-MATB-6792.

### Western blots

Total protein was quantified using a BCA protein assay kit (Pierce). 10 μg of protein was loaded per well, and samples were separated on 12% SDS-PAGE gels in Tris-glycine running buffer (25 mM Tris, 192 mM glycine, 0.1% SDS). Proteins were then electroblotted onto PVDF membrane (40V, 3 hrs) in transfer buffer (25 mM Tris-HCl, 192 mM glycine, 20% methanol), the membrane blocked in TBS-T (50 mM Tris-HCl, 150 mM NaCl, 0.1% Tween 20) with 5% non–fat milk powder for 1 hour. The membrane was incubated with primary antibodies overnight at 4° with the following dilutions: anti-NDUFS1 (rabbit monoclonal, ab169540, 1:30,000), anti-NDUFS3 (mouse monoclonal, 0.15 mg/ml, ab110246), or anti-alpha tubulin loading control (mouse monoclonal, ab7291, 1:30,000). Following incubation with the primary antibodies, blots were washed 3 × 5 min, and 2 × 15 mins in TBS-T and incubated with the appropriate secondary antibody for 1 hour at room temperature: goat anti-rabbit HRP (Invitrogen) 1:30,0000; goat anti-mouse HRP (Dako) 1:3,000. After secondary antibody incubations, blots were washed and proteins of interested detected using ECL prime chemiluminescent detection reagent (GE Healthcare) and the blots imaged using an ImageQuant LAS 4000 (GE Healthcare). Signals were normalised to the loading control using ImageJ [78].

### Oxidative phosphorylation enzyme assays and oxygen consumption

Cell lysates were prepared 48 hours post-transfection, by harvesting cells by trypsinisation, washing three times in ice cold phosphate buffered saline followed by centrifugation to pellet the cells (2 mins, 1000 g). Cell pellets were resuspended to homogeneity in KME buffer (100 mM KCl, 50 mM MOPS, 0.5 mM EGTA, pH 7.4) and protein concentrations were determined using a BCA protein assay detection kit (Pierce). Cell lysates were flash frozen in liquid nitrogen, and freeze-thawed three times prior to assay. 300-500 μg of cell lysate was added per assay, and assays were normalised to the total amount of protein added.

All assays were performed using a Shimadzu UV-1800 spectrophotometer, absorbance readings were taken every second and all samples were measured in duplicate. Activity of complex I (CI, NADH:ubiquinone oxidoreductase) was determined by measuring the oxidation of NADH to NAD^+^ at 340 nm at 30° in an assay mixture containing 25 mM potassium phosphate buffer (pH 7.2), 5 mM MgCl_2_, 2.5 mg/ml fatty acid free albumin, 0.13 mM NADH, 65 μM coenzyme Q and 2 μg/ml antimycin A. The decrease in absorbance was measured for 3 mins, after which rotenone was added to a final concentration of 10 μM and the absorbance measured for a further 2 mins. The specific complex I rate was calculated as the rotenone-sensitive rate minus the rotenone-insensitive rate. Complex II (CII, succinate dehydrogenase) activity was determined by measuring the oxidation of DCPIP at 600 nm at 30°. Lysates were added to an assay mixture containing 25 mM potassium phosphate buffer (pH 7.2) and 2 mM sodium succinate and incubated at 30° for 10 mins, after which the following components were added, 2 μg/ml antimycin A, 2 μg/ml rotenone, 50 μM DCPIP and the decrease in absorbance was measured for 2 mins. Complex III (CIII, Ubiquinol:cytochrome *c* oxidoreductase) activity was determined by measuring the oxidation of decylubiquinol, with cytochrome *c* as the electron acceptor at 550nm. The assay cuvettes contained 25 mM potassium phosphate buffer (pH 7.2), 3 mM sodium azide, 10 mM rotenone and 50 μM oxidized cytochrome *c*. Decylubiquinol was synthesized by acidifying decylubiquinone (10mM) with HCl (6M) and reducing the quinine with sodium borohydride. After the addition of 35 μM decylubiquinol, the increase in absorbance was measured for 2 mins. Activity of Complex IV (CIV, cytochrome *c* oxidase) was measured by monitoring the oxidation of cytochrome *c* at 550 nm, 30° for 3 min. A 0.83 mM solution of reduced cytochrome *c* was prepared by dissolving 100 mg of cytochrome *c* in 10 ml of potassium phosphate buffer, and adding sodium ascorbate to a final concentration of 5 mM. The resulting solution was added into SnakeSkin dialysis tubing (7 kDa molecular weight cutoff, Thermo Scientific) and dialyzed against potassium phosphate buffer, with three changes, at 4° for 24 hrs. The redox state of the cytochrome *c* was assessed by evaluating the absorbance spectra from 500-600 nm. The assay buffer contained 25 mM potassium phosphate buffer (pH 7.0) and 50 μM reduced cytochrome *c*. The decrease in absorbance at 550 nm was recorded for 3 mins. As a control the enzymatic activity of the tricarboxylic acid cycle enzyme, citrate synthase (CS) was assayed at 412 nm at 30°C in a buffer containing 100 mM Tris-HCl (pH 8.0), 100 μM DTNB (5,5-dithiobis[2-nitrobenzoic acid]), 50 μM acetyl coenzyme A, 0.1% (w/v) Triton X-100 and 250 μM oxaloacetate. The increase in absorbance was monitored for 2 mins.

The following extinction coefficients were applied: complex I (CI), ε = 6220 M^−1^ cm^−1^, CII, ε = 21,000 M^−1^ cm^−1^; CIII, ε = 19,100 M^−1^ cm^−1^; CIV, ε = 21,840 M^−1^ cm^−1^ (the difference between reduced and oxidised cytochrome *c* at 550 nm); CS, ε = 13,600 mM^−1^ cm^−1^.

Oxygen consumption was measured using an extracellular oxygen sensitive probe (MitoXpress Xtra, Luxcel Biosciences). Cells were plated at 50,000 cells/well in a 96 well plate in complete media and allowed to adhere overnight. Cells were then assayed by addition of MitoXpress Xtra, and oxygen consumption assessed by time-resolved fluorescence (380 nm excitation, 650nm emission; integration window 1, 30 μs delay, 30 μs gate time; integration window 2, 70 μs delay, 30 μs gate time).

### Oxidative stress measurements

Hydrogen peroxide production was assessed as a marker of reactive oxygen species generation using the fluorescent indicator Amplex Red (10 μM, Invitrogen) in combination with horseradish peroxidise (0.1 units ml^−1^). Total amount of H_2_O_2_ produced was normalised to mg of protein added. Protein carbonylation was assessed using the OxyBlot protein oxidation detection kit (Merck Millipore), and differential carbonylation was assessed by densitometry. The cell stress assay was performed on cells seeded in 48 well plates, and assayed 12 hours later by the addition of (final concentration): rotenone (5 μM), malonate (40 μM), antimycin A (500 μM), oligomycin (500 μM), sodium azide (3 mM), NaCl (300 mM), CaCl_2_ (5.4 mM). Cells were heat shocked at 42° and UV irradiated using a Stratlinker UV Crosslinker for 10 minutes (2.4 J cm^−2^). Cell viability was assessed by the addition of Alamar Blue (Invitrogen) according to the manufacturer’s instructions.

## Acknowledgements

This work was funded by the Wellcome Trust (CPP, TMS, OBR, SRG), a European Research Council Advanced Grant (CPP, ACM, TMS, KR), the Medical Research Council (CPP, WH), a Marie Curie Intra-European Career Development Award (ACM), the University of Oxford (ACM), the Royal Society (ACM), the Clarendon Fund (JYT), the Natural Sciences Engineering Research Council of Canada (JYT), and Diabetes UK (LCH, 11/0004175). Microarray hybridizations were performed by the OXION array facility. We would like to thank Andrew Dodd for critical reading of the manuscript.

## Author contributions

TMS, ACM and CPP conceived the study. TMS, KR, SRG and NL performed all experiments. WH performed computational analyses (FANTOM5 and GTEx data). ACM made initial MRE predictions and OBR made updated predictions. SC provided *Dicer* knock-out embryonic stem cells, experimental protocols, aid and advice in the culturing of these cells, JYT helped conceive the ES cell experiments and assisted with ES cell culture. LH provided OXPHOS experimental protocols and assisted with assays and interpretation. CPP supervised the study. TMS and CPP wrote the manuscript.

